# ZC3H11A interacts with PABPN1 and alters polyadenylation of viral transcripts

**DOI:** 10.1101/2023.03.09.531914

**Authors:** Katharina Kases, Erik Schubert, Zamaneh Hajikhezri, Mårten Larsson, Leif Andersson, Göran Akusjärvi, Tanel Punga, Shady Younis

## Abstract

Nuclear mRNA metabolism is regulated by multiple proteins, which either directly bind to RNA or form multi-protein complexes. The RNA-binding protein ZC3H11A is involved in nuclear mRNA export, and NF-κB signaling and is essential during mouse embryo development. Furthermore, previous studies have shown that ZC3H11A is important for nuclear-replicating viruses. However, detailed biochemical characterization of the ZC3H11A protein has been lacking. In this study, we established the ZC3H11A protein interactome in human and mouse cells. We demonstrate that the nuclear poly(A)-binding protein PABPN1 interacts specifically with the ZC3H11A protein and controls ZC3H11A localization into nuclear speckles. We report that ZC3H11A specifically interacts with the human adenovirus type 5 (HAdV-5) capsid mRNA in a PABPN1-dependent manner. Notably, ZC3H11A uses the same zinc finger motifs to interact with PABPN1 and mRNA, with a single zinc finger motif responsible for these functions. Further, we demonstrate that the lack of ZC3H11A alters the polyadenylation of HAdV-5 capsid mRNA. Taken together, our results suggest that the ZC3H11A protein acts as a novel regulator of polyadenylation of nuclear mRNA.

## Introduction

RNA-binding proteins (RBPs) control multiple steps in gene expression, including gene transcription, RNA export, RNA stability, and mRNA translation (1, 2). The RBPs directly interact with the target mRNAs and make multiple contacts with other proteins to achieve temporal and spatial regulation of gene expression. ZC3H11A is an RNA-binding protein with three uncharacterized CCCH-type zinc finger motifs (ZFM) motifs present at the N-terminus of the protein (3). Several studies have shown that ZC3H11A is a subunit of the Transcription-Export (TREX) complex, which is responsible for the nuclear mRNA export (4–9). TREX is a multiprotein complex containing the hexameric core THO complex (THOC1, THOC2, THOC3, THOC5, THOC6, THOC7), which in turn associates with several subunits including ZC3H11A (8). Essential for TREX-mediated mRNA export is its interaction with properly processed mRNAs (10). Therefore, different TREX complex subunits associate with the mRNA 5’ cap structure (NCBP1, NCBP3), exon junctions (DDX39A/B, ALYREF(THOC4), UIF, LUZP4, CHTOP), and the 3’ end poly(A) tail (THOC5, ZC3H14) (8, 11). Further, the TREX complex stimulates recruitment of the m^6^A reader protein YTHDC1 to mRNA and favors exporting m^6^A-modified mRNA (12). Together, these interactions ensure that capped, spliced, polyadenylated, and m^6^A-modified mRNAs are recognized and efficiently exported from the nucleus. In addition to mRNA recognition, the TREX complex interacts with the nuclear pore complex (NPC) to accomplish mRNA export to the cytoplasm. This interaction is mediated by the TREX subunit TAP/NXF1 as it interacts with the nucleoporins at NPC and mRNA along with the ALYREF and THOC5 subunits (13, 14).

Although the exact biochemical characteristics of ZC3H11A have remained enigmatic, it has been characterized as a heat-shock-induced protein controlling virus growth (15–17). Four different human nuclear-replicating viruses (Human Immunodeficiency Virus 1 (HIV-1), Human Adenovirus 5 (HAdV-5), Influenza A virus (IAV), and Herpes Simplex Virus 1 (HSV-1)) showed reduced growth in HeLa cells lacking the ZC3H11A protein (16). Detailed studies in HAdV-5-infected cells revealed that the ZC3H11A protein was relocalized to replication centers in HAdV-5-infected HeLa cells and that a lack of ZC3H11A reduced cytoplasmic accumulation of some viral mRNAs, suggesting its role in virus mRNA processing and nuclear export (16). Similarly, a lack of the ZC3H11A protein affected porcine pseudorabies virus (PRV) and porcine circovirus 2 (PCV2) amplification by reducing the cytoplasmic accumulation of viral mRNA in porcine cells (15). Collectively, the ZC3H11A protein is essential for virus replication, probably due to its role in TREX complex-mediated virus mRNA export (3, 15). Two very recent studies have further shed light on ZC3H11A functions. First, it has been shown that ZC3H11A is crucial for the viability and survival of mouse embryos (9). Notably, genetic characterization of *Zc3h11a^-/-^* mice embryos revealed dysregulation of the fatty acid and glycolysis pathways (9). Second, the human ZC3H11A protein can suppress NF-κB signaling, thereby controlling the expression of several interferon-stimulated genes in the HeLa cells (18).

The poly(A)-binding proteins (PABPs) are the proteins that specifically bind to polyadenosine (poly(A)) tails present at the 3’ end of mRNA. The PABPs are grouped based on their subcellular localization: i) nuclear PABPs (PABPNs: PABPN1 and PABPN1L), and ii) cytoplasmic PABPs (PABPCs: PABPC1, PABPC3, PABPC4, PABPC4L, PABPC5, and ePAB) (19). The PABPs lack catalytic domains but instead function as scaffold proteins as they bind to poly(A) tails via their RNA recognition motifs (RRMs) (20). The best-known function of PABPN1 is to stimulate poly(A) polymerase (PAP)-dependent polyadenylation (21, 22). In addition, PABPN1 can also define the poly(A) tail length as it mediates interactions between the PAP protein and the cleavage and specificity factor 1 (CPSF1) (22, 23). Historically, the PABPN1 protein binding to poly(A) tails was suggested to increase mRNA stability, nuclear mRNA export, and translation (24, 25). Surprisingly, PABPN1 also has an opposite role as it can promote the turnover of some of the non-coding snoRNA host gene (SNHG) transcripts (26–28). Available data suggest that PABPN1 is a subunit of a poly(A) tail exosome targeting (PAXT) complex, which can target polyadenylated SNHG transcripts for degradation (27).

While ZC3H11A is essential for mouse development (9) and for the growth of multiple nuclear-replicating viruses (15, 16), the detailed biochemical characterization of the protein has been lacking. Therefore, the present study aimed to reveal the biochemical characteristics of the human ZC3H11A protein and correlate them to its functions.

## Results

### ZC3H11A interacts with the TREX complex in human and mouse cells

Our recent study established the ZC3H11A interactome in mouse embryonic stem cells (mESCs) (9). To understand whether the human ZC3H11A protein interactome differs from the mouse counterpart, we performed mass spectrometry (MS)-based proteomics analysis and compared the ZC3H11A protein interacting partners in human HeLa cells and mESCs as outlined in Figure 1A. The results of HeLa and mESCs MS analyses showed a significant interaction between ZC3H11A and proteins involved in mRNA processing, nuclear export, and mRNA 3’ end processing (Figure 1B and Supplementary Figure 1). The most significant ZC3H11A-interacting partners both in the mESCs and HeLa cells were the TREX complex subunits (the THOC proteins, ALYREF, CHTOP, POLDIP3) and PABPN1 (Figure 1C). Whereas it is known that ZC3H11A copurifies with the TREX complex subunits, its interaction with the PABPN1 protein has not been characterized previously.

**Figure 1.**
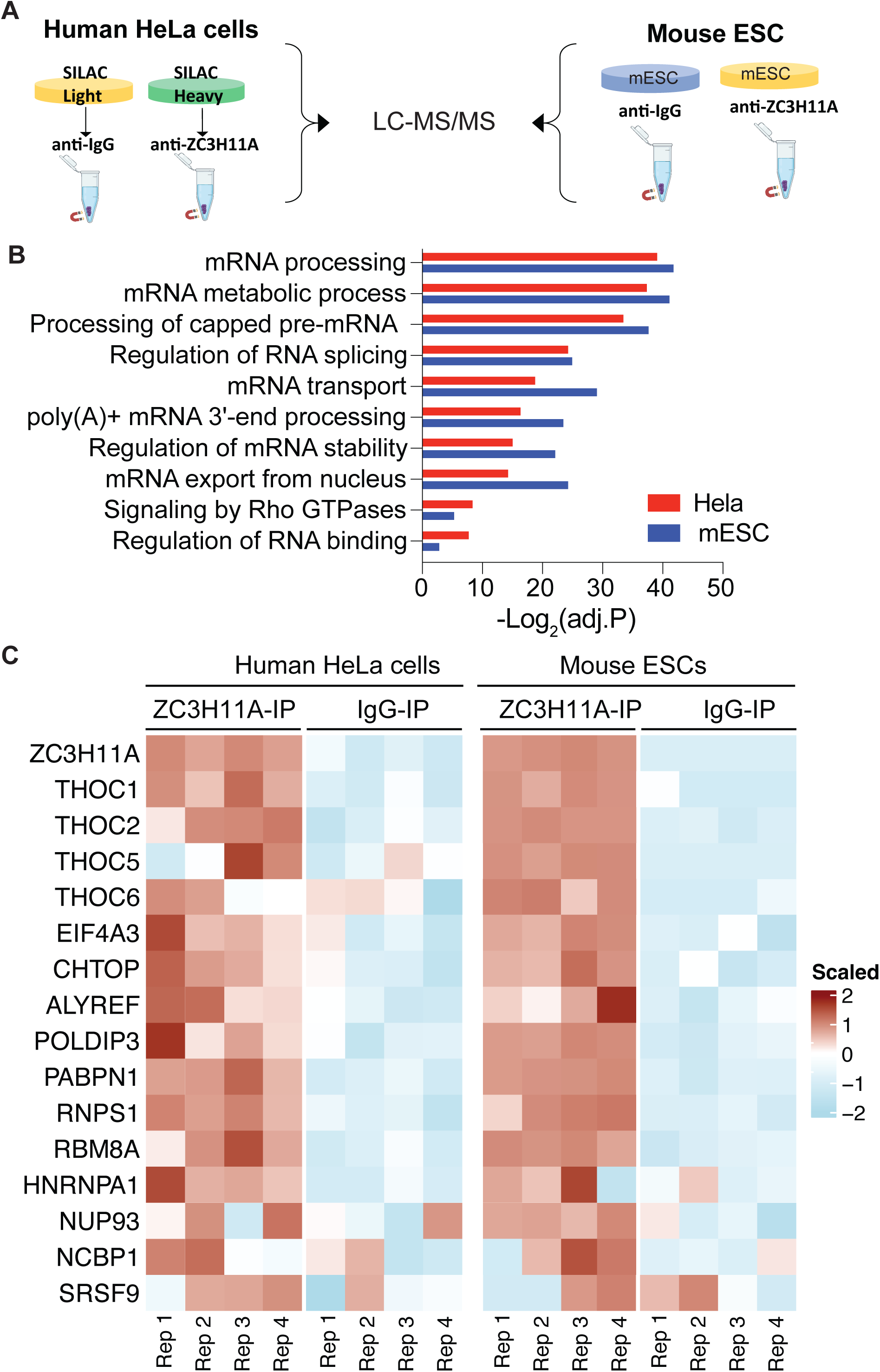
A comparison between the human and mouse ZC3H11A interactome. **(A)** Schematic illustration of the co-immunoprecipitation (co-IP) mass spectrometry (MS)-based experiment using the anti-ZC3H11A and anti-IgG antibodies in human HeLa cells and mouse embryonic stem cells (mESCs). **(B)** A pathway analysis of the identified interacting partners with endogenous ZC3H11A in HeLa cells and mESC. (**C)** Heatmap of ZC3H11A-interacting partners involved in RNA export and 3’ end processing, based on ZC3H11A co-IPs in HeLa cells and mESCs. The heatmap represents the scaled values of log2-intensity of the indicated proteins (adjusted *P* <0.05).

### The ZC3H11A zinc finger motifs mediate interactions with the PABPN1 protein

Our previous study showed that deletion of the three ZC3H11A zinc finger motifs (ZFMs, deletion referred to as dZF) affected ZC3H11A functions (16). Therefore, to further reveal the details of the human ZC3H11A interactome, we overexpressed the ZC3H11A(wt), and the ZC3H11A(dZF) proteins in parallel and affinity purified the proteins to identify interacting partners by MS, as depicted in Figure 2A. In line with its proposed mRNA export function (6, 8–10, 14), the GFP-ZC3H11A(wt) protein specifically interacted with several of the known TREX complex subunits (THOC1, THOC2, THOC6, DDX39B, CHTOP, POLDIP3, RBM8A, and ALYREF) and again with the PABPN1 protein (Figure 2B). Notably, the interaction between ZC3H11A and PABPN1 was significantly reduced by the deletion of the ZFMs (Figure 2C). On the other hand, interactions between ZC3H11A and some of the TREX complex proteins THOC2, POLDIP3, and ALYREF were not affected by the same deletion (Figure 2D). To confirm that our results were not biased due to the N-terminal GFP-tag, we overexpressed and immunopurified the ZC3H11A(wt)-Flag protein in HeLa cells and analyzed the interacting proteins by MS. Similarly to the GFP- ZC3H11A(wt), the ZC3H11A(wt)-Flag protein interacted with the PABPN1 protein (Supplementary Figure 2). Our MS-based findings encouraged us to investigate how the individual ZFMs contribute to the ZC3H11A protein function and its interaction with the PABPN1 proteins.

**Figure 2.**
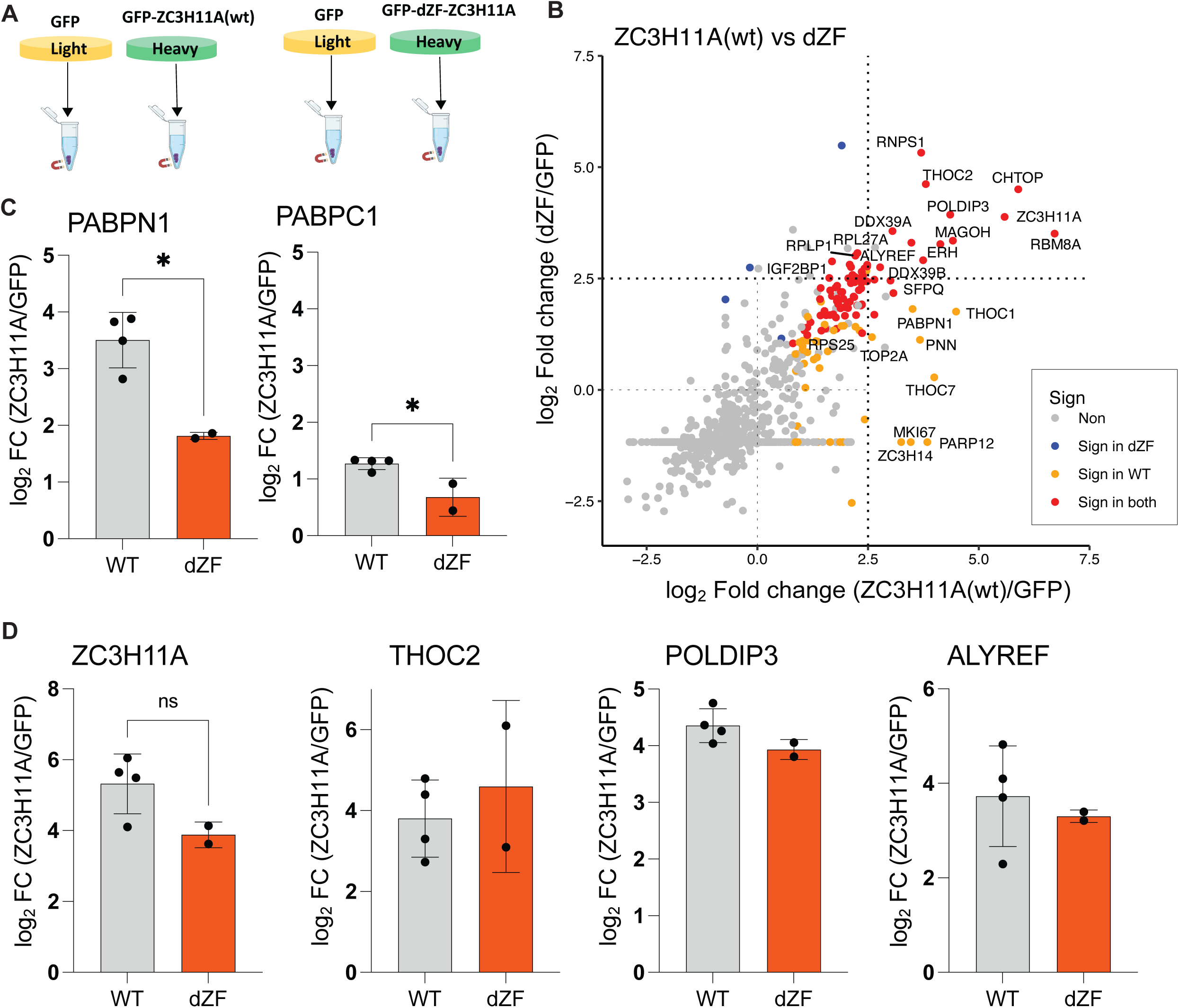
Human ZC3H11A protein interacts with the TREX complex subunits and PABPs. **(A)** Schematic illustration of the co-IP experiments using GFP, GFP- ZC3H11A(wt), and GFP-ZC3H11A(dZF) as the baits in HeLa cells. **(B)** Scatter plot showing enrichment of co-IP proteins from ZC3H11A(wt)/GFP vs. ZC3H11A(dZF)/GFP proteins in HeLa cells. No: not significant; Sign: significant at adj *P* <0.05 in the indicated condition. **(C)** Log2 enrichment of the PABPN1 and PABPC1 proteins interacting with the ZC3H11A(wt) or ZC3H11A(dZF) proteins. **(D)** Log2 enrichment of the ZC3H11A, THOC2, POLDIP3 and ALYREF proteins detected in the ZC3H11A(wt) vs. ZC3H11A(dZF) co-IPs. * corresponds to adjusted *P* <0.05, ns: not significant.

### Individual ZFMs control ZC3H11A binding to PABPN1

To better understand the human ZC3H11A protein functions, we focused on its interaction with the PABPN1 protein as both proteins localize into nuclear speckles and impact the mRNA nuclear export (16, 29, 30). The ZC3H11A protein contains three CCCH-type ZFMs at the N-terminus of the protein (Figure 3A). Since the functions of single ZFMs have not been studied, we mutated individual ZFMs by replacing the first cysteine in the CCCH motif with an alanine (CCCH>ACCH). This type of mutation reduces Zn^2+^-binding by disrupting the formation of a functional ZFM (31). First, we tested whether the triple cysteine mutation (C8A/C37A/C66A, referred to as 3C>A) alters ZC3H11A binding to PABPN1. Indeed, the 3C>A mutation blocked ZC3H11A interaction with PABPN1 (Figure 3B). Also, a deletion mutant ZC3H11A(dZF) was deficient in binding to PABPN1. To further delineate the role of ZFMs, we tested how individual ZFM mutations affect ZC3H11A binding to PABPN1. Notably, mutations within the first (C8A) and third (C66A) ZFM eliminated binding, whereas the C37A mutation reduced ZC3H11A binding to PABPN1 (Figure 3C). ZC3H11A and PABPN1 are mRNA-binding proteins; hence their mutual interaction might be mediated by RNA. However, as shown in Figure 3D, the ZC3H11A- PABPN1 interaction was not altered in the cell extracts treated with ribonuclease A (RNase A), indicating that ZC3H11A interacts with PABPN1 in an RNA-independent manner. Since ZC3H11A interacts with the THOC proteins (Figure 1C) (6, 8), we tested if the ZFM mutations also affected ZC3H11A binding to two THOC proteins: THOC1 and THOC5. Both tested proteins interacted with both the wild-type and 3C>A mutant proteins (Figures 3E and 3F), confirming our MS-based analysis (Figure 2D) that some of the THO complex members interact with ZC3H11A independently of the ZFMs.

**Figure 3.**
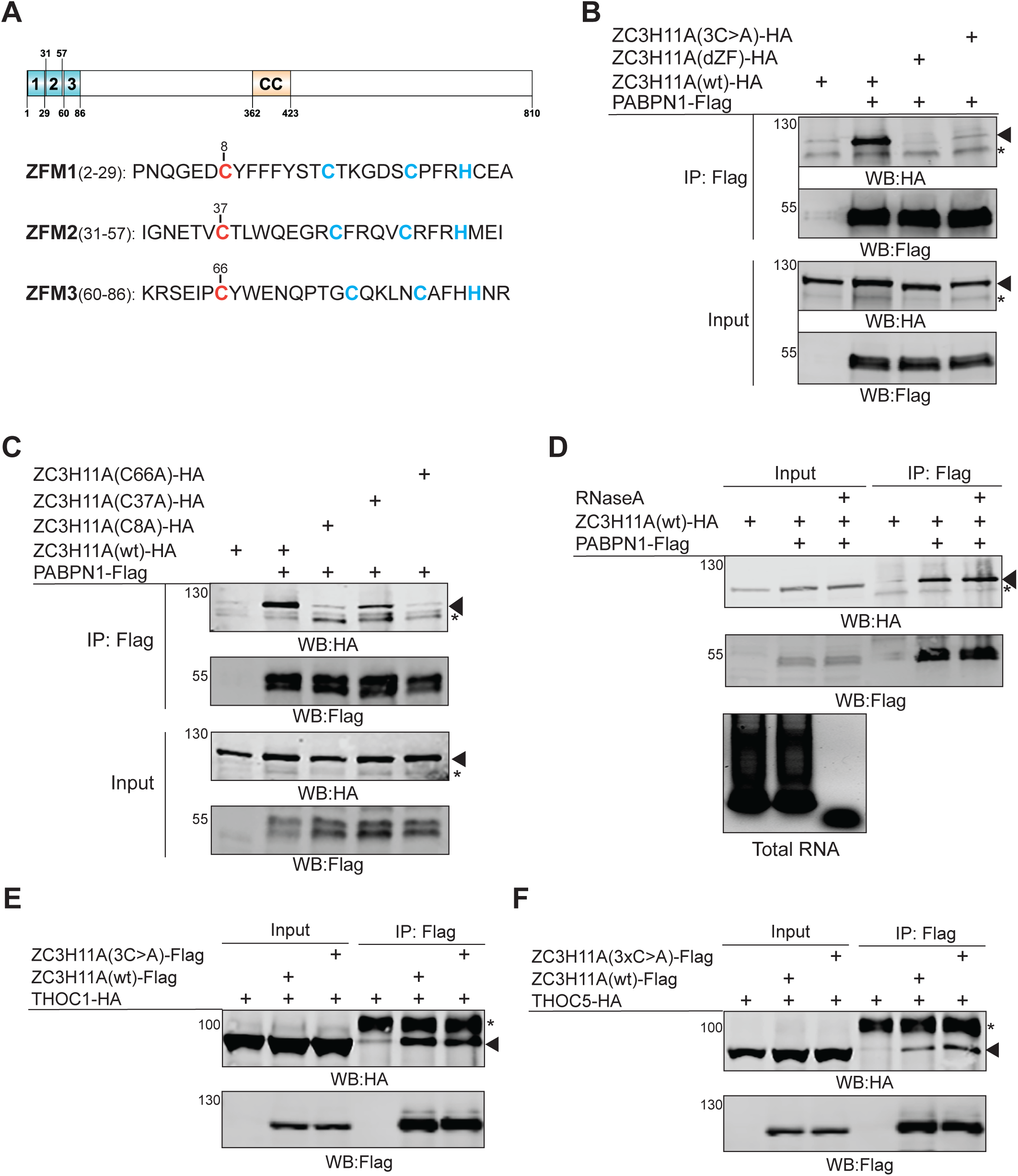
Individual ZFMs control ZC3H11A binding to PABPN1. **(A)** A schematic drawing of the human ZC3H11A protein. Individual zinc finger motifs (ZFM1-3) and coil-coiled domain (CC) are indicated based on the uniport.org (O75152) annotation. Individual CCCH motifs are shown (red and blue). Mutated cysteine (Cys(C) to Ala(A)) is indicated in red. **(B)** Co-IP of the HA-tagged ZC3H11A proteins using PABPN1-Flag as the bait in transfected HEK293T cells. The 3C>A mutant includes C8A, C37A, and C66A triple mutations, whereas the dZF mutant lacks the entire N-terminus of the protein. Western blot (WB) analysis with the anti-Flag and anti-HA antibodies. An asterisk indicates an unspecific protein; arrowheads indicates migration of the ZC3H11A protein. **(C)** Co-IP of the individual ZC3H11A ZFM mutants with the PABPN1-Flag protein. **(D)** Co-IP of ZC3H11A(wt)-HA with PABPN1- Flag as a bait in the absence or presence of RNase A treatment. The efficiency of RNase A treatment in the cell lysates is shown on an agarose gel image below the protein images. **(E)** Co-IP of THOC1-HA with either ZC3H11A(wt)-Flag or ZC3H11A(3C>A)-Flag as the baits. **(F)** Co-IP of THOC5-HA with either ZC3H11A(wt)-Flag or ZC3H11A(3C>A)-Flag as the baits. An asterisk indicates an unspecific protein, whereas the arrowheads indicate migration of the THOC1-HA and THOC5-HA proteins.

Collectively, our data indicate that human ZC3H11A specifically associates with PABPN1 via ZFMs and that the first and third ZFM are essential for this binding.

### A single ZFM is needed for ZC3H11A localization into nuclear speckles

ZC3H11A and PABPN1 are nuclear proteins with characteristic localization in the nuclear speckles (30). Nuclear speckles (NS) are distinct nuclear structures containing the proteins involved in mRNA processing and export (30). Therefore, we analyzed whether individual ZFM mutations alter ZC3H11A localization into NS. To this end, we performed indirect immunofluorescence experiments in HeLa cells transiently expressing the ZC3H11A-Flag proteins, followed by detecting both endogenous PABPN1 and transfected ZC3H11A-Flag proteins. Similarly to the previous study (16), the ZC3H11A protein was detected in distinct NS, which overlapped PABPN1 staining (Figure 4). Analysis of different ZFM mutants revealed that the individual ZFMs mutations, C3>A, and dZF mutations expelled the ZC3H11A protein from the NS. Interestingly, all the tested mutant proteins showed strong nuclear staining, suggesting that ZFM mutations affect only localization to NS but not to the nucleus. Overall, our microscopy experiments align with the co-immunoprecipitation experiments (Figure 3), showing that the integrity of the individual ZFMs is needed to target ZC3H11A into the NS and mediate its binding to the PABPN1 protein.

**Figure 4.**
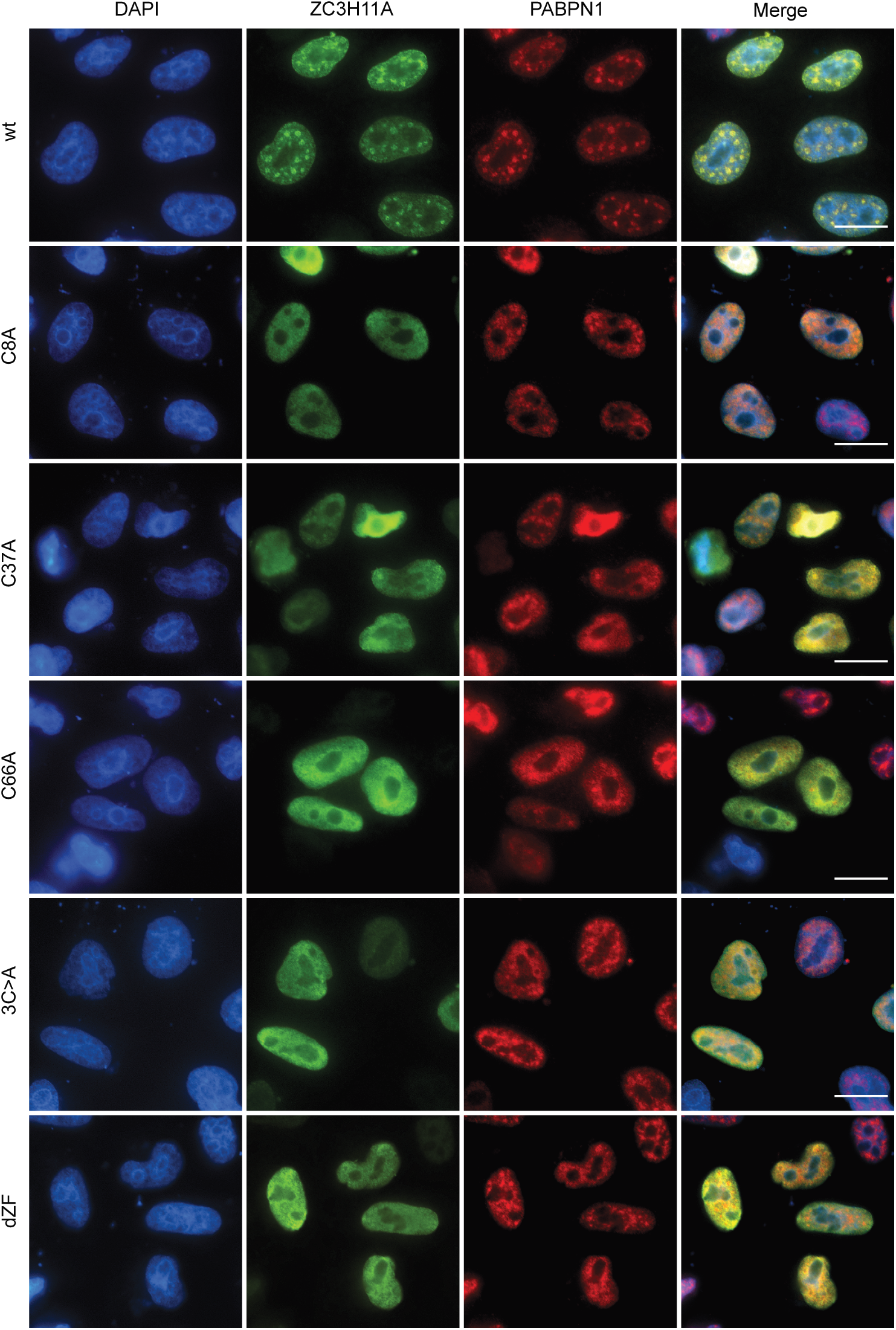
Individual ZFMs define ZC3H11A localization into nuclear speckles. HeLa cells were transfected with the respective ZC3H11A-Flag encoding plasmids for 24 h. Indirect immunofluorescence (IF) was done with the anti-Flag and anti- PABPN1 antibodies. Cells were counterstained with DAPI. Scalebar is 20 μm.

### PABPN1 retains ZC3H11A in the nuclear speckles

Since our data indicate that ZC3H11A interacts with PABPN1 (Figure 3) and that this interaction occurs in the NS (Figure 4), we hypothesized that PABPN1 is needed for the ZC3H11A protein accumulation in the NS. To test this, we treated HeLa cells with siRNA against PABPN1 (siPABPN1) and analyzed the subcellular distribution of the ZC3H11A(wt)-Flag and ZC3H11A(dZF)-Flag proteins by indirect immunofluorescence. Notably, siPABPN1 treatment excluded the ZC3H11A(wt) protein from the NS into diffused nuclear staining (Figure 5A). Remarkably, the diffused nuclear staining pattern of the ZC3H11A(wt)-Flag protein in siPABPN1- treated cells (Figure 5A, the second-row images) resembled the PABPN1-binding deficient ZC3H11A(dZF)-Flag protein staining pattern in the siScr-treated cells (Figure 5A, the third row images). To understand if the siPABPN1-treatment had a general effect on NS formation, the siPABPN1-treated cells were stained with an antibody detecting a known NS protein, the splicing factor SC35(SRSF2) (32). However, as shown in Figure 5B, the lack of PABPN1 did not detectably alter the SC35 protein localization.

**Figure 5.**
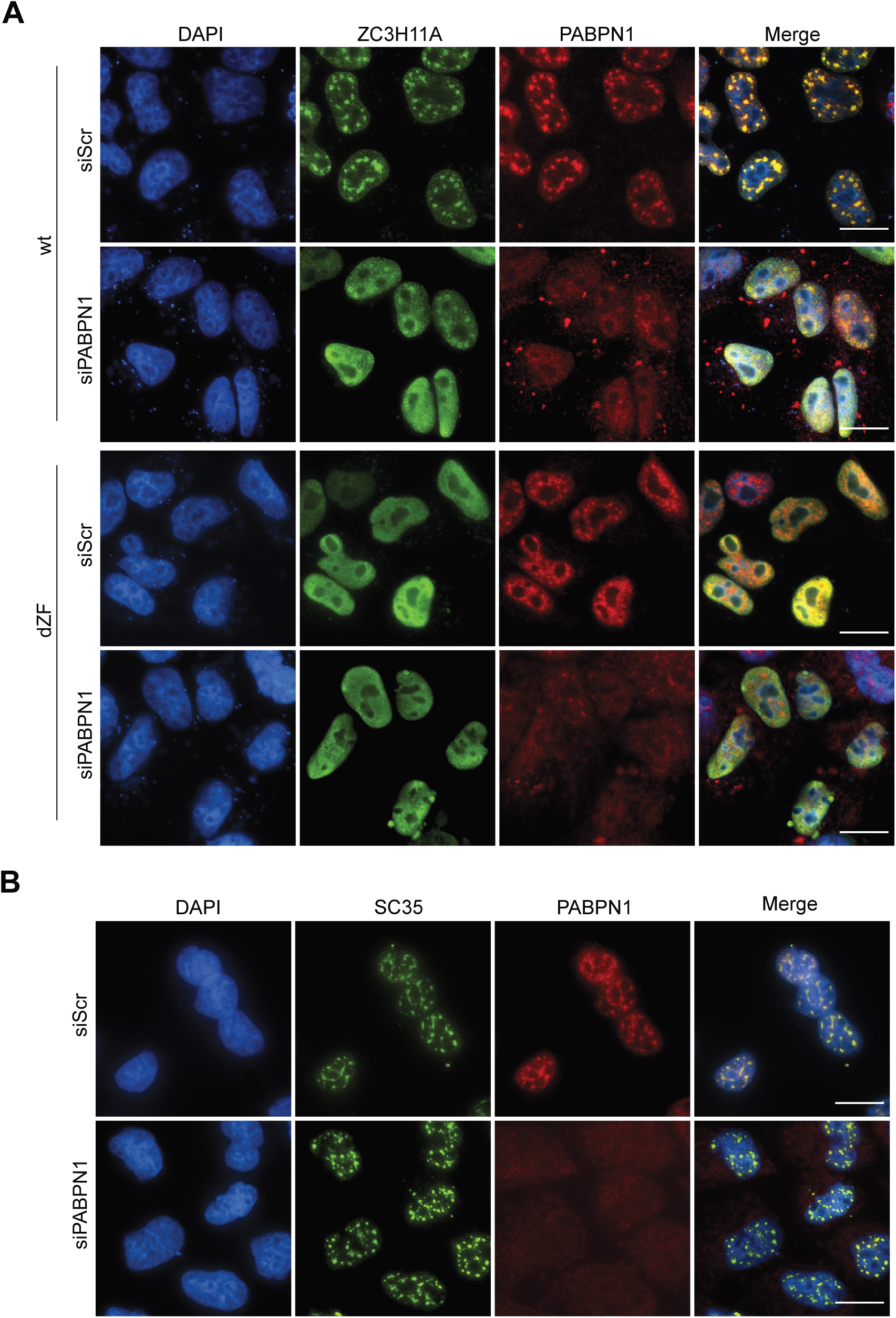
PABPN1 is needed for ZC3H11A localization in nuclear speckles. **(A)** HeLa cells were treated with scrambled (siScR) or PABPN1-specific (siPABPN1) siRNAs (24 h) followed by transient expression of the ZC3H11A(wt)-Flag or ZC3H11A(dZF)-Flag proteins (24 h). IF was done using the anti-Flag and anti- PABPN1 antibodies. **(B)** HeLa cells were treated with siScR or siPABPN1, followed by the detection of the SC35 and PABPN1 proteins by IF. Cells were counterstained with DAPI. Scalebar is 20 μm.

Taken together, our data support the hypothesis that PABPN1 is needed for the localization of ZC3H11A into NS in HeLa cells.

### The ZC3H11A protein contains two independent nuclear localization signals

Since the ZC3H11A protein shows intense nuclear staining, we were interested to know whether ZC3H11A contains a nuclear localization signal (NLS) driving its nuclear localization. The classical NLS (cNLS) is defined as a short, 4-8 amino acid peptide sequence enriched with positively charged amino acids such as lysines (K) and arginines (R) (33). Bioinformatic analysis using the cNLS Mapper program revealed that the ZC3H11A protein contains a potential NLS between amino acids 284-304 (34). Since this region contains three K and two R residues, we generated a ZC3H11A mutant protein (KR5>A), where all five residues were exchanged for alanines. The KR5>A mutant protein was expressed and detected with immunofluorescence in HeLa cells. Surprisingly, the KR5>A mutant protein showed a very similar nuclear localization as did the ZC3H11A(wt)-Flag protein (Figure 6A), suggesting that additional NLS may exist in the protein. This was also supported by the observation that the C-terminal deletion mutant ZC3H11A(1-280) showed apparent cytoplasmic staining, whereas mutant 1-530 was still found in the nucleus.

**Figure 6.**
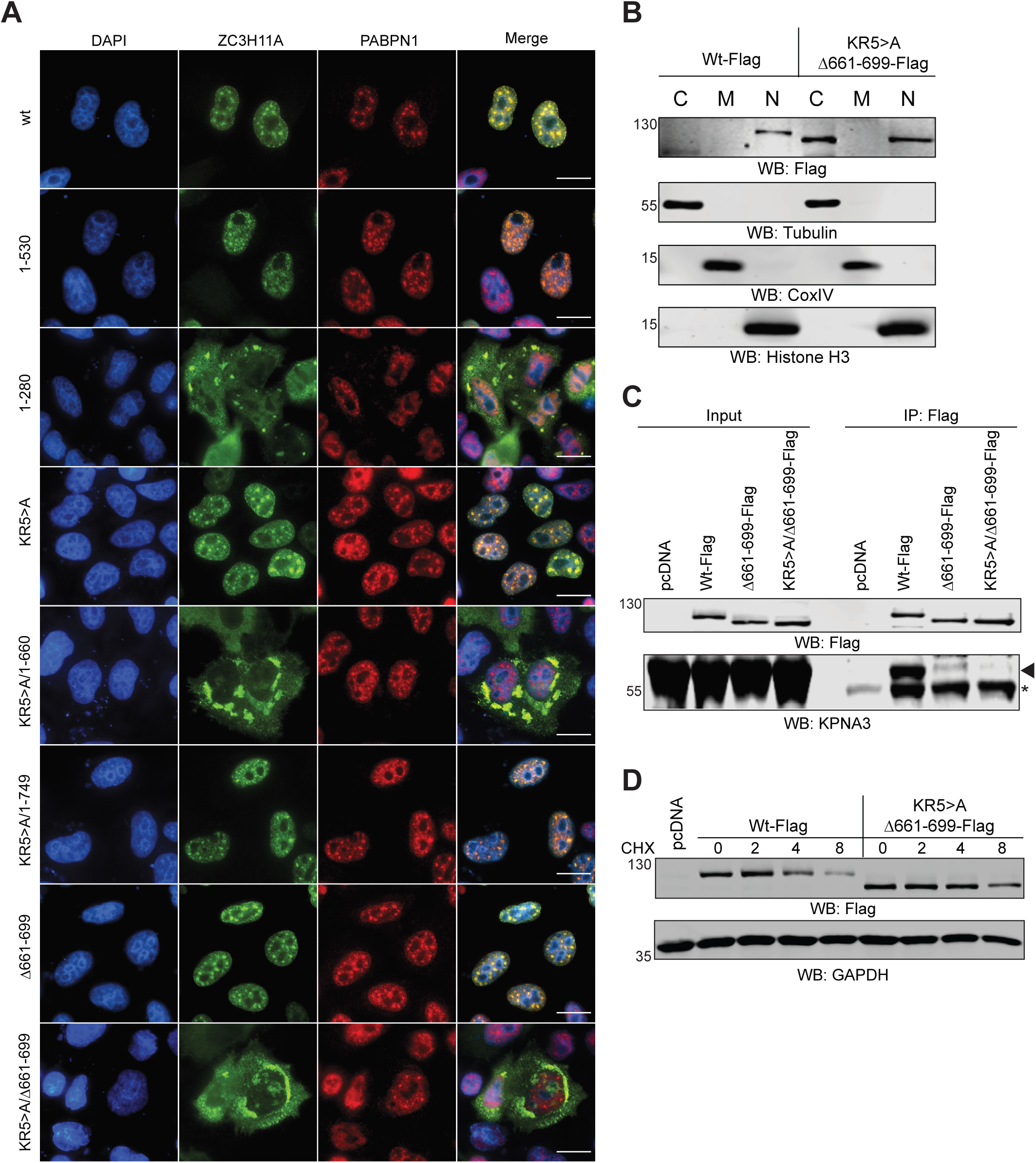
The ZC3H11A protein contains two independent nuclear localization signals. **(A)** Subcellular localization of the respective ZC3H11A-Flag proteins. IF was done using the anti-Flag and anti-PABPN1 antibodies; cells were counterstained with DAPI. Scalebar is 20 μm. **(B)** Biochemical fractionation of the ZC3H11A(wt)-Flag and NLS-deficient ZC3H11A(KR5>A/Δ661-699)-Flag proteins. Western Blot analysis with the anti-Flag, anti-tubulin (marker for cytoplasmic fraction), anti-CoxIV (a marker for mitochondrial fraction), and anti-histone H3 (marker for nuclear fraction). **(C)** Co- IP of the endogenous KPNA3 using the ZC3H11A(wt)-Flag, ZC3H11A(Δ661-699)- Flag, and ZC3H11A(KR5>A/Δ661-699)-Flag proteins as the baits in HeLa cells. WB analysis with the anti-Flag and anti-KPNA3 antibodies. An asterisk indicates an unspecific protein, whereas the arrowheads indicates the migration of the KPNA3 protein. **(D)** Cycloheximide (CHX) treatment of HeLa cells expressing the ZC3H11A(wt)-Flag or ZC3H11A(KR5>A/Δ661-699)-Flag proteins. Cells were collected at 0, 2, 4, and 8 h after CHX was added to the growth media.

Therefore, we generated two C-terminal deletion mutants using the ZC3H11A(KR5>A) sequence as the background. As shown in Figure 6A, deletion mutant KR5>A/1-660 showed intense cytoplasmatic staining, whereas the mutant KR5>A/1-749 was detected only in the nucleus. To further narrow potential NLS, we generated a deletion mutant protein lacking amino acids 661-699. The ZC3H11A(Δ661-699)-Flag protein still showed apparent NS staining, whereas the double mutant ZC3H11A(KR5>A/Δ661-699) was excluded from the nucleus. Hence, the ZC3H11A protein contains at least two independent NLS: one between amino acids 284-304 and another between amino acids 661-699. This is also supported by our biochemical fractionation experiment where the ZC3H11A(wt)-Flag protein was detected only in the nucleus, whereas the double NLS mutant protein ZC3H11A(KR5>A/Δ661-699) was found in the cytoplasm and to some extent still in the nucleus (Figure 6B).

If the ZC3H11A protein contains functional NLS, it should specifically interact with the importin receptors responsible for binding to NLS and nuclear delivery of the proteins (35). Therefore, we tested whether a known importin receptor subunit α-3 (KPNA3), which copurifies with the TREX complex (5), can bind to the ZC3H11A protein. The ZC3H11A(wt)-Flag specifically interacted with the endogenous KPNA3, whereas the NLS-deficient proteins either showed drastically reduced binding (Δ661- 699) or lacked the binding (KR5>A/Δ661-699) (Figure 6C). In our previous study we showed that ZC3H11A is a relatively unstable protein (16). Therefore, we tested whether ZC3H11A subcellular localization influences its stability by treating ZC3H11A(wt)-Flag or ZC3H11A(KR5>A/Δ661-699)-Flag expressing HeLa cells with cycloheximide (CHX). The CHX treatment blocks *de novo* protein synthesis and therefore enables the detection of protein decay during a predefined time (36). Notably, the ZC3H11A(KR5>A/Δ661-699) protein was more resistant to degradation compared to the wild-type counterpart (Figure 6D).

Taken together, our results suggest that the ZC3H11A protein contains two independent NLS, which are needed for nuclear localization and fast protein turnover.

### Lack of ZC3H11A alters mRNA polyadenylation

Our previous study showed that ZC3H11A inactivation reduced HAdV-5 lytic growth in HeLa cells (16). Notably, the lack of ZC3H11A altered viral capsid mRNA export in this study. Since polyadenylation stimulates the mRNA export (37), we hypothesized that the ZC3H11A protein might interfere with HAdV-5 capsid mRNA polyadenylation. The HAdV-5 capsid precursor mRNA (pre-mRNA) is processed into five different mRNA families (L1 to L5) due to the usage of different polyadenylation sites (L1pA site to L5pA site) (Figure 7A). To test our hypothesis, we used the extension Poly(A) Test (ePAT), which is a PCR-based method where PCR amplicons reflect the actual length distribution of the poly(A) tail on the target mRNA (38). As the amplification control, a TVN-PAT reaction was used. Due to an anchored PCR primer, the TVN-PAT amplicon has an invariant poly(A)_12_ tail, irrespective of the sample’s actual poly(A)-lengths (38). To test the effect of ZC3H11A and PAPBPN1 on polyadenylation, we used siRNAs to knock down protein expression (Figure 7B). As an additional control, cells were treated with a siRNA targeting the Cleavage and Polyadenylation Factor 1 (CPSF1), which is a subunit of the essential CPSF complex recognizing the poly(A) signal on the pre-mRNA (39). Analysis of the TVN-PAT amplicons revealed very similar PCR amplification at L3 and L5 pA sites in siScR- and siZC3H11A-treated samples (Figure 7C). As expected, siCPSF1-treatment blocked PCR amplification, hence validating the specificity of the method. Also, the siPABPN1-treatment reduced the TVN-PAT signal, although not to the same extent as observed for siCPSF1. Interestingly, the ePAT reactions showed less intense (L3 and L5) and slower migrating (L3) amplicons in the siZC3H11A-treated samples (Figure 7C). In line with the reduced TVN-PAT signal, the ePAT signal was almost undetectable in siPABPN1- and siCPSF1-treated cells. This indicates that lack of ZC3H11A alters the polyadenylation of viral transcripts using the L3 (i.e., pVI, hexon) and L5 (i.e., fiber) poly(A) sites (Figures 7A and 7C). In addition to its mRNA stabilizing role (39), the PABPN1 protein has been shown to cause SNHG degradation as it is a subunit of the exonucleolytic PAXT complex (27). Therefore, we tested whether ZC3H11A alters the accumulation of PAXT-targeted SNHG10 and SNHG19 RNA (27). As shown in Figure 7D, siZC3H11A treatment did not change SNHG10 and SNHG19 expression, whereas siPABPN1-treatment elevated the levels of these two SNHGs in HeLa cells.

**Figure 7.**
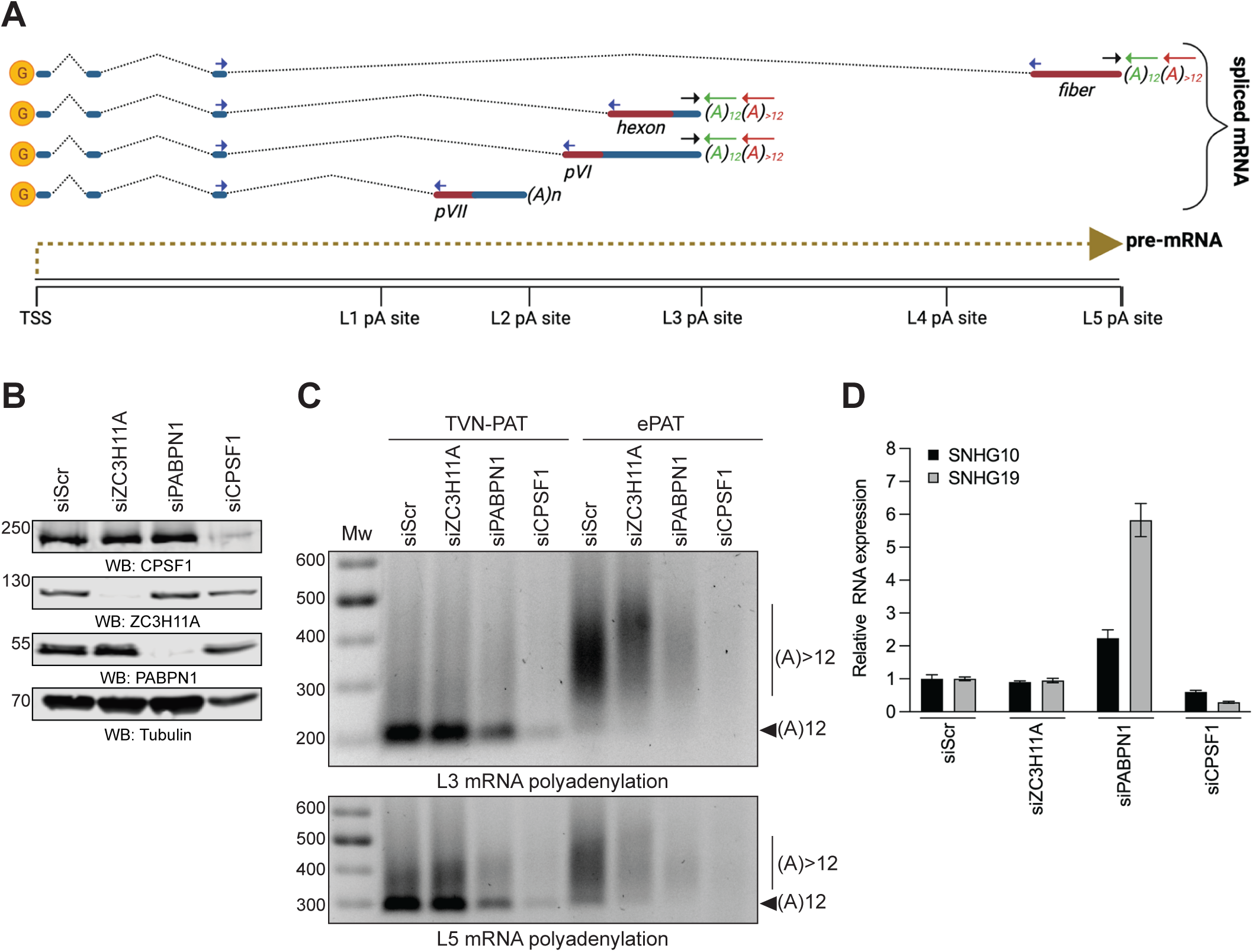
ZC3H11A deficiency alters HAdV-5 mRNA polyadenylation. **(A)** Simplified drawing of some HAdV-5 capsid mRNAs (pVII, pVI, hexon, fiber) originating from the virus major late transcription unit (MLTU). Blue arrows indicate spliced mRNA amplicon, a combination of black and green arrows indicate TVN-PAT amplicon, a combination of black and red arrows indicates ePAT amplicon. TSS; transcription start site, pA; polyadenylation site, G; capped nucleotide. **(B)** Western blot from siRNA-treated HeLa cells. Proteins were detected with the anti-ZC3H11A, anti-PABPN1, anti-CPSF1, and anti-tubulin antibodies. **(C)** Total RNA from siRNA-treated and HAdV-5-infected samples was analyzed with ePAT and TVN-PAT methods. PCR amplicons detect HAdV-5 L3 and L5 poly(A) tails. An arrow with the label “(A)12” indicates the migration of mRNA with defined 12 As containing poly(A) tail. A vertical line with the label “(A)>12” indicates mRNA species with poly(A) tail longer than 12 As. **(D)** Expression of SNHG10 and SNHG19 RNA in siRNA-treated HeLa cells. HPRT1 was used to normalize SNHG expression. Data are shown as mean +/- SD from a triplicate experiment.

Collectively, our data indicate that the ZC3H11A protein alters HAdV-5 capsid mRNA polyadenylation but is not engaged in cellular SNHG10 or SNHG19 RNA degradation.

### ZC3H11A binds to HAdV-5 mRNA in a PABPN1-dependent manner

Since ZC3H11A ZFMs interact with both polyadenylated mRNA (16) and the PABPN1 protein (Figure 3), we hypothesized that PABPN1 might mediate ZC3H11A binding to mRNA. To test this, we analyzed ZC3H11A binding to some of the HAdV-5 capsid mRNAs (pVII, hexon, fiber) (Figure 8) in the lung epithelial cell line A549, a standard model cell line of many respiratory viruses, including HAdV-5 (40). First, we tested how different ZC3H11A ZFM mutations influence ZC3H11A binding to viral mRNAs. As shown in Figure 8A, all ZFM mutants (C8A, C37A, C66A, C3>A) showed reduced binding to the tested viral transcripts. Interestingly, RNA-binding of the mutant C8A was most severely affected compared to other ZFM mutant proteins. Since the same ZC3H11A mutant proteins were deficient in binding to PABPN1 (Figure 3), we hypothesized that the PABPN1 protein might mediate ZC3H11A binding to viral mRNAs. In agreement with this hypothesis, siPABPN1-treatment blocked ZC3H11A binding to two tested viral transcripts (Figure 8B). The ZC3H11A(wt)-Flag protein was immunopurified equally well from the siScr- and siPABPN1-treated cells (inserted panel in Figure 8B), indicating that the reduced RNA-binding was not caused by a lower level of the ZC3H11A(wt)-Flag protein in the siPABPN1-treated cells (inserted panel in Figure 8B). Proper processing of the capsid mRNAs is essential for forming infectious HAdV-5 particles. Therefore, we assumed that a lack of ZC3H11A should affect the production of infectious progeny virus in A549 cells. To test this, siZC3H11A- and siPABPN1-treated A549 cells were infected with a HAdV-5 virus expressing the NanoLuc (NLuc) reporter gene. Cell lysates were prepared 24 hours post-infection and used to re-infect HeLa cells, followed by NLuc measurement as a read-out of the formation of infectious virus particles. Both siZC3H11A- and PABPN1-treatments affected NLuc expression, with a more drastic effect in siPABPN1-treated cells (Figure 8C). To further confirm these observations, we generated an A549 cell line (ZC3^KO^) where the ZC3H11A protein expression was eliminated using the CRISPR/Cas9 editing approach. As shown in Figure 8D, NLuc reporter gene expression was reduced in HeLa cells re-infected with ZC3^KO^ cell lysates.

**Figure 8.**
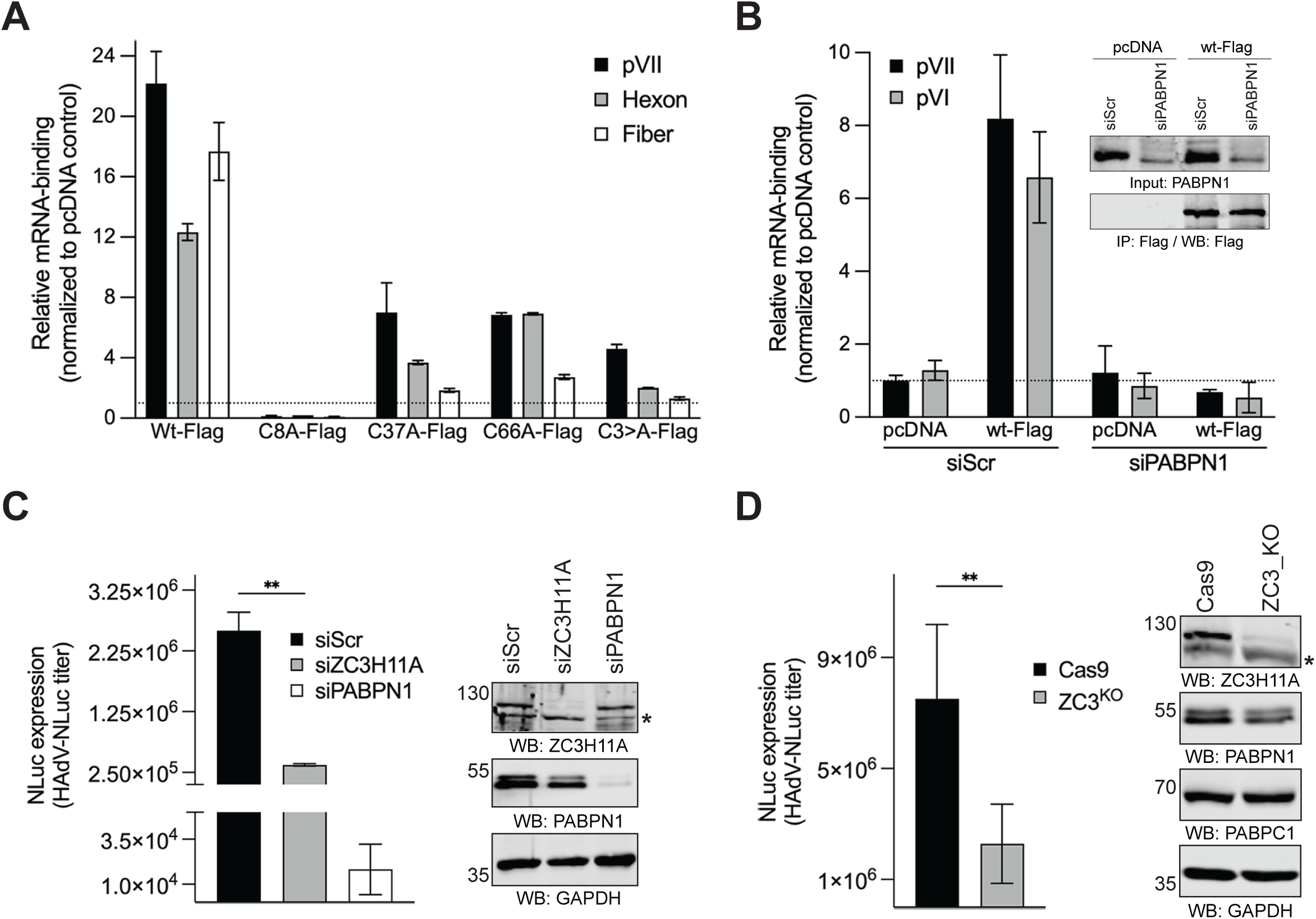
ZC3H11A binds to mRNA in a PABPN1-dependent manner. **(A)** Individual ZC3H11A-Flag proteins were immunoprecipitated from HAdV-5-infected (MOI=5, 48 h) A549 cells and analyzed for binding to target mRNA (pVII, Hexon, and Fiber) using qRT-PCR. Data are shown relative mRNA enrichment in the ZC3H11A- Flag plasmid transfected samples compared to the control pcDNA plasmid transfected sample. Mean and +/- SD from a triplicate experiment are shown. **(B)** PABPN1 is needed for the ZC3H11A-Flag protein binding to mRNAs. The ZC3H11A(wt)-Flag protein was immunoprecipitated from siScr- or siPABPN1-treated (48 h) HAdV-5-infected A549 cells (MOI=5, 24 h) and analyzed for viral mRNA (pVII and pVI) binding. Mean and +/- SD from a triplicate experiment are shown. Insert shows the equal recovery of the ZC3H11A(wt)-Flag protein from siScr- and siPABPN1-treated and HAdV-5-infected cells. **(C)** Formation of infectious progeny (HAdV-NLuc titer) in siRNA-treated (48 h) A549 cells. Expression of the NanoLuc (NLuc) reporter gene was used as the read-out of infectious virus particles. NLuc values are shown as mean +/- SD from a triplicate experiment. Western blot shows the efficiency of the siZC3H11A- and siPABPN1-treatment (48 h) in A549 cells. **(D)** Formation of infectious progeny in CRISPR/Cas9-modified A549 cells. Control (Cas9) and ZC3H11A modified (ZC3_KO) A549 cells were infected and analyzed as in panel C. Western blot showing lack of the ZC3H11A protein in CRISPR/Cas9 modified A549 cells. An unpaired t-test indicated a significant reduction in NLuc expression in ZC3H11A-lacking cells (**, *P*<0.005).

## Discussion

Most ZC3H11A functional studies reported so far are based on siRNA- or CRISPR/Cas9-based elimination of the ZC3H11A protein. In the present study, we reveal the human ZC3H11A protein interactome with a clear focus on characterizing the ZC3H11A-PABPN1 interaction.

The most significant ZC3H11A interacting proteins identified in this study were those involved in the mRNA processing, nuclear mRNA export, and mRNA 3’ end processing (Figure 1B). ZC3H11A interaction with the mRNA export proteins was expected as it is known to copurify with the TREX complex members (4–6, 8, 9). Notably, the ZC3H11A protein has been shown to interact with the TREX core protein THOC2, TREX subunits POLDIP3 and UAP56(DDX39B) in an ATP-dependent and RNA-independent manner (7). The observations that lack of ZC3H11A causes nuclear accumulation of polyadenylated mRNAs and that ZC3H11A interacts with the members of the TREX complex have put forward a model whereby the ZC3H11A protein regulates nuclear mRNA export (3, 7, 16). Surprisingly, the ZC3H11A protein also interacted with the PABPN1 protein, implying that it might be involved in mRNA 3’ end processing in addition to targeting the TREX complex and mRNA export. In line with our report, a previous study identified ZC3H11A as one of the PABPN1 interacting proteins due to DNA damage response (DDR). However, the functional consequences of this interaction during DDR were not elucidated (41). Notably, we show that the PABPN1 protein interacts with the ZC3H11A ZFMs, which allowed us to dissect the functional consequences of the ZC3H11A-PABPN1 interaction (Figure 3).

ZC3H11A is an RNA-binding protein, and deletion of all three ZFMs abolished its binding to polyadenylated mRNAs (16). Further, based on the resemblance to other CCCH-type RNA-binding proteins (e.g., ZAP or ZFP36) (3), it is predicted that the ZC3H11A ZFMs mediate its binding to mRNA. Therefore, it was surprising that PABPN1 interacted with the same protein region needed for the ZC3H11A RNA-binding activity. Further, our protein-protein interaction and protein-RNA binding experiments revealed that the mutation within the first ZFM (C8A) most efficiently blocked ZC3H11A interaction with PABPN1 and HAdV-5 mRNA (Figures 3 and 8). Although mutations within the other two ZFMs (C37A and C66A) also affected interaction with PABPN1 and viral mRNAs, the effects seen with the first ZFM (C8A) were the most drastic. This suggests that the first ZFM has a pivotal role in ZC3H11A interaction with PABPN1 and mRNA. Since the same C8A mutation affected ZC3H11A binding to PABPN1 and viral mRNAs, we hypothesized that the PABPN1 protein mediates ZC3H11A binding to mRNA as it interacts with the poly(A) tail. We have previously characterized the role of the ZC3H11A protein in HAdV-5 mRNA metabolism in the HeLa cells (16); therefore we continued to use this model system to reveal the details of the ZC3H11A-PABPN1-mRNA interaction. Indeed, in PABPN1-depleted cells, the ZC3H11A protein binding to viral mRNAs was abolished (Figure 8B). The PABPN1 depletion did not detectably affect the transfected ZC3H11A(wt)-Flag protein levels, indicating that ZC3H11A reduced RNA-binding is caused by the lack of the PABPN1 protein on targeted mRNAs. Since PABPN1 binds to the poly(A) tails, it implies that the ZC3H11A-RNA interactions should occur preferably at the 3’ end of mRNA. Indeed, a clear enrichment of the ZC3H11A protein binding at the 3’ end of both cellular and viral mRNAs has been shown (16). Similarly, the preferential binding of the ZC3H11A protein at the 3’ end of cellular mRNAs was reported in a study analyzing RNA-binding of 150 different RBPs (42). The same study also indicated that the ZC3H11A might be needed to export polyadenylated mRNAs. Together, our data and data from other studies suggest that the PABPN1 protein mediates ZC3H11A RNA-binding at the 3’ end of mRNA.

Since the PABPN1 protein controls the poly(A) tail synthesis and length (25), we also investigated if the ZC3H11A protein has an impact on poly(A) tail status using HAdV-5 as a model. Since reliable detection of the poly(A) tails is challenging, we used the PCR-based ePAT detection method, which can detect qualitative changes in the poly(A) tail (38). Indeed, our data show that the ePAT method is reliable as inhibition of pre-mRNA cleavage (siCPSF1-treatment) or poly(A) synthesis (siPABPN1-treatment) blocked detection of the oligoadenylated ((A_12_), TVN-PAT) and polyadenylated ((A_>12_), ePAT) viral L3 and L5 mRNAs (Figure 7B). Interestingly, the absence of ZC3H11A did not detectably interfere with the oligoadenylation (A_12_) of the L3 and L5 mRNAs but instead altered their polyadenylation (A_>12_). More specifically, siZC3H11A treatment reduced L3 mRNA polyadenylation, whereas surprisingly, the length of the poly(A) tail was increased on the L5 mRNAs. These observations suggest that ZC3H11A acts specifically at the polyadenylation step ((A_>12_) and is probably finetuning the length of the poly(A) tail, depending on the origin of the mRNA. Since the ZC3H11A protein is not involved in PABPN1- dependent SNHG transcript turnover (Figure 7D), we believe the ZC3H11A-PABPN1 interaction is operational only in the canonical mRNA polyadenylation process. This is also supported by the fact that ZC3H11A is not part of the PAXT complex, which can target polyadenylated SNHG transcripts for degradation (27). In our previous study, we showed that the lack of the ZC3H11A protein caused nuclear retention of HAdV-5 mRNA in HeLa cells (16). It is reasonable to assume that one of the underlying reasons for that is altered polyadenylation of the viral transcripts, which will be retained in the cell nucleus. Hence, the data from our present and previous study (16) indicate that ZC3H11A might control HAdV-5 mRNA polyadenylation and export of viral capsid mRNAs in HeLa cells.

Our study also reveals the details of the ZC3H11A protein subcellular localization. The ZC3H11A protein is known to localize into NS (16, 43), and based on our study, this localization pattern is controlled by the individual ZFMs (Figure 4). All three tested ZFM mutations (C8A, C37A, C66A) blocked ZC3H11A co-localization with PABPN1 in NS. Further, after siPABPN1-treatment ZC3H11A failed to associate with the NS (Figure 5). These observations indicate that PABPN1 is essential to keep ZC3H11A in the NS. This particular localization pattern raises a question of what is the function of the ZC3H11A-PABPN1 complex in the NS. The NS are proposed to regulate gene expression and RNA processing. This is supported by several MS studies showing that NS contain multiple splicing factors and proteins involved in mRNA polyadenylation (e.g., CPSF1, PABPN1) (30, 44). Therefore, the ZC3H11A- PABPN1 interaction in the NS might either be needed to mark particular mRNA species for mRNA export or to control their poly(A) tail length.

Another important finding of our study was the identification of two NLSs in ZC3H11A. Since the ZFM mutations or deletions did not expel the ZC3H11A protein from the nucleus, it was evident that the protein must contain strong NLS, which will drive its nuclear localization. Our two identified sequence elements fulfill the criteria for NLSs as both contain the basic amino acids (K and R). The N-terminal NLS, which was predicted to locate between aa 284-304, can be regarded as a bipartite NLS (BP-NLS) with the first 3 basic aa (K286, R287, and K288) separated from the last two basic aa (K300 and R301) by a 12 aa spacer sequence. Hence, the ZC3H11A BP-NLS resembles the BP-NLS previously identified in the 53BP1 and ING4 proteins (33). We did not do mutational analysis within the C-terminal NLS (aa 661-699), although the presence of the tripeptide motif KRK (aa 669-671) would suggest the presence of a monopartite NLS. Remarkably, only mutations of both NLS elements (KR5>A/Δ661-699) blocked ZC3H11A-KPNA3 interaction efficiently, which coincided with predominant cytoplasmic localization of ZC3H11A (Figures 6A and 6C). However, our biochemical fractionation revealed that the ZC3H11A(KR5>A/Δ661-699) still localized into the nucleus to some extent (Figure 6B), indicating the possibility for an additional, uncharacterized NLS. Hence, our mutational analysis shows that two essential structural elements are needed for ZC3H11A to function as a nuclear protein. First, the integrity of the ZFMs is required for PABPN1-mediated ZC3H11A localization into the NS and binding to HAdV-5 mRNA. Second, the general localization of ZC3H11A into the cell nucleus is driven by the monopartite and bipartite NLSs and their interaction with the KPNA3 protein.

The ZC3H11A protein has a fast turnover rate, although the mechanistic details about its proteasomal degradation have not been revealed (16). Notably, the cytoplasmic ZC3H11A (KR5>A/Δ661-699) was more stable compared to the wild-type protein (Figure 6D), indicating that ZC3H11A might undergo proteasomal degradation in the nucleus. Indeed, a recent study has suggested that NS might act as the hotspots of the nuclear ubiquitination (45).

Collectively, our study reveals human and mouse ZC3H11A interactomes and shows that the ZC3H11A protein specifically targets the PABPN1 protein and mRNA polyadenylation machinery.

## Experimental procedures

### Cell lines

A549 cells were obtained from ATCC. HEK293T and HeLa cells were obtained from Dr. Johan Eriksson, Uppsala University, Sweden, and Dr. Thomas Dobner, Leibniz Institute of Virology, Germany, respectively. The cell lines were grown in Dulbecco’s modified Eagle medium (DMEM, Thermo Fisher Scientific) supplemented with 10% fetal bovine serum (FBS, ThermoFisher) and penicillin-streptomycin solution (PEST, Thermo Fisher Scientific) at 37°C in a 5% CO_2_ incubator. The mouse embryonic stem cells (mESCs) were obtained as a gift from Dr. Alice Jouneau, Université Paris-Saclay, France. mESCs were cultured on gelatin-coated plates, and maintained in DMEM supplemented with 10% heat-inactivated fetal bovine serum, penicillin (0.2 U/mL), streptomycin (0.2 µg/mL), L-glutamine (0.2 µg/mL) (Gibco) and with recombinant mouse Leukemia Inhibitory Factor (LIF, 20 U/ml, Millipore).

### Generation of an A549-KO cell line

CRISPR/Cas9-edited A549 cells (ZC3_KO) were created as described previously (16). Briefly, a DNA fragment containing the U6 promoter, the gRNA sequence, the gRNA scaffold sequence, and a termination signal (IDT) was co-transfected to A549 cells with a Cas9-expressing plasmid and a linear hygromycin marker (Clontech). Transfected cells were cultured in selective media with 100 μg/ml of hygromycin B for 2 weeks. Control A549 cells (Cas9) were transfected only with the Cas9-expressing plasmid and a linear hygromycin marker. The resistant single cell clones were isolated and screened for efficient ZC3H11A knockout by Western blotting.

### Transfection, plasmids, and siRNA

All plasmid transfections were performed with JetPrime (Polyplus) transfection reagent according to the manufacturer’s protocol. Plasmids expressing the PABPN1- Flag (NM_004643) and THOC5-HA (NM_001002878.1) proteins were purchased from GenScript. Plasmid pcDNA-THOC1-HA was generated by cloning the THOC1 sequence from plasmid phHpr1-GST (Addgene #11200) into pcDNA3(C-HA) background using BamHI/XhoI restriction enzymes. Plasmids expressing the ZC3H11A proteins were generated by cloning the ZC3H11A cDNA from pAcGFP1C1-ZC3H11A plasmid (16) into pcDNA3(C-HA) or pcDNA3(C-Flag) vectors using EcoRI/XhoI restriction enzymes. The expressed ZC3H11A proteins contain C-terminal HA- or Flag- antibody epitope tags. Point mutations C8A, C37A, C66A, 3C>A (contains C8A, C37A, and C66A triple mutation), KR5>A (contains K286A, R287A, K288A, K300A, and R301A mutations) were done using QuikChange Mutagenesis Kit (Agilent). Deletion mutants (1-280, 1-530, 1-660, and 1-749) were generated using PCR primers annealing to the respective sequences. All three ZFM deletion mutant (ZC3H11A(dZF)) was generated by cloning the corresponding cDNA sequence (amino acids 87-810) from pAcGFP1C1-ZC3H11A(ZF^Δ3^) (16) into the pcDNA3(C-HA) and pcDNA3(C-Flag) plasmids. Mutation Δ661-699 was performed by cloning the ZC3H11A gBlock gene fragment (IDT) lacking the corresponding sequence into the pcDNA3-ZC3H11A(wt)-Flag vector using the PflMI and XhoI restriction enzymes. The siScr (5’-AGGUAGUGUAAUCGCCUUG-3’), siPABPN1 (5’- GUAGAGAAGCAGAUGAAUAdTdT-3’) and siCPSF1 (5’-GCAUUUCGCUGCUGCGCUA-3’) were purchased from Eurofins Genomics. siGENOME human ZC3H11A siRNA (M-021238-01-0005, Horizon Discovery) was used to knock down ZC3H11A mRNA. All siRNAs transfections (25 nM, 48 h) were done using JetPrime reagent.

### Virus infection

Virus infections were done using replication-competent HAdV-5 as described previously (46). Replication-competent HAdV-5(NLuc) virus (generously provided by ZedCe Medicals AB, Sweden) was engineered by inserting the PGK-NLuc-poly(A) cassette from pNL1.1.PGK (Promega) plasmid into virus genome. Infections were performed at a multiplicity of infection (MOI) of 5, defined as fluorescence-forming units (FFU/cell), in infection media (DMEM + 2% FBS without any other supplements). After 1 h incubation at 37°C in a 5% CO_2_ incubator, virus-containing infection media was replaced with the normal growth media (DMEM containing FBS and PEST).

### Co-immunoprecipitation, subcellular fractionation, and western blotting

HEK293T cells were grown on 6-well plates and transfected with the respective plasmids (2 µg of plasmid/well) for 36 h. Cells were harvested and lysed in 350 μl of lysis buffer (50mM Tris-HCl pH7.5, 150mM NaCl, 0.5% NP-40, 0.5% Triton X-100, 0.1% sodium deoxycholate (SDC), 0.2μM ZnCl_2_) supplemented with Halt protease inhibitors (Thermo Fisher Scientific) for 30 min on ice. The soluble cell lysate was used for immunoprecipitation with an anti-Flag(M2) affinity gel (A2220, Sigma) overnight at 4°C. The beads were washed 2x1ml in lysis buffer and 2x1 ml of lysis buffer containing 300 mM NaCl. RNase A treatments were done by incubating cell lysates in the presence of RNase A (25 ng/μl as final concentration) for 20 min at 37°C. Total RNA was isolated from RNase A-treated or non-treated cell lysates (ca. 20% of the whole lysate) using TRIzol LS reagent (Thermo Fischer Scientific) according to the manufacturer’s protocol. Isolated RNA was separated on a 1% agarose gel (1XTBE) and visualized using GelRed dye (Biotium). Both inputs (corresponds to 5%) and immunoprecipitates were separated on 9% SDS-PAGE, transferred to a nitrocellulose membrane, and detected with the anti-Flag (rabbit, Sigma, F1804 or rabbit Proteintech, 20543-1-AP), anti-HA (mouse, Biolegend) and anti-KPNA3 (rabbit, ABclonal, A8347) antibodies. Proteins were visualized with fluorescence-labeled secondary antibodies IRDye® 680 or IRDye® 800 (LI-COR) using the Odyssey CLX imaging system (LI-COR).

Subcellular fractionation of HeLa cells transfected with the ZC3H11A-Flag expressing plasmids (5μg plasmid/100mm tissue culture plate) was done using Cell Fractionation Kit (Abcam, ab109719) according to the manufacturer’s instructions. The anti-histone H3 (rabbit, Abcam, ab1791), anti-tubulin (mouse, Santa Cruz, sc- 69969), and anti-Cox IV (rabbit, Proteintech, 11242-1-AP) antibodies were used to validate the purity of the nuclear, cytoplasmic and mitochondrial fractions, respectively. Whole-cell lysates were prepared as described previously (47), and proteins were detected using the anti-PABPN1 (rabbit, Abcam, ab75855), anti- ZC3H11A (rabbit, Abcam, ab99930), anti-CPSF1 (mouse, Santa Cruz, sc-166281), anti-GAPDH (mouse, Proteintech, 60004-1-Ig), anti-PABP (rabbit, Abcam, ab21060), and anti-actin (goat, Santa Cruz, c-1616) antibodies. Plasmid transfected HeLa cells were treated with cycloheximide (CHX, Sigma, C4859, 100 μg/ml as final concentration) 24 h post-transfection. Cells were collected 2, 4, and 8 h after the start of treatment, lysed and analyzed by western blotting as described previously (36).

### Co-immunoprecipitation and mass spectrometry sample preparation

HeLa cells were cultured in DMEM for Stable Isotope Labeling with Amino acids in Cell culture (SILAC, Thermo Fisher Scientific) supplemented with 10% dialyzed FBS (MWCO 10 kDa, Thermo Fisher Scientific), 100 U/ml of penicillin (Thermo Fisher Scientific), 100 μg/ml of streptomycin (Thermo Fisher Scientific), 0.25 μg/ml of amphotericin B (Thermo Fisher Scientific) and light isotopic labels L -arginine-HCl and L -lysine-2 HCl or heavy isotopic labels ^13^C_6_, _15_N4 L-arginine-HCl (Arg-10) and ^13^C_6_, ^15^N_2_ L-lysine-2 HCl (Lys-8) (Thermo Fisher Scientific). To avoid contamination of light amino acids, sub-culturing was performed using Cell dissociation buffer (Thermo Fisher Scientific) instead of trypsin. Isotopic incorporation was checked using a script in R as previously described (48) after approximately five cell divisions to assess complete (>95%) labeling. Conversion of arginine to proline was checked by calculating the percentage of heavy proline (Pro-6) containing peptides among all identified peptides, and kept at <5%. Labeled HeLa cells (light or heavy) were grown on 100 mm tissue plates, and total protein lysates were prepared using Pierce IP lysis buffer (Thermo Fisher Scientific) supplemented with protease inhibitors (Complete Ultra Tablets, Roche) and Pierce Universal Nuclease (Thermo Fisher Scientific). Lysates were cleared by centrifugation at 20,000 x *g* for 10 min at 4°C, and rotated end-over-end at 4°C with anti-IgG (Abcam, light lysate), and anti- ZC3H11A (Atlas Antibodies, heavy lysate) antibodies in Protein LoBind tubes (Eppendorf). Thereafter, 30 µl of Dynabeads Protein G (Thermo Fisher Scientific) was added to each tube and incubated for 30 min at room temperature, followed by washing three times with Pierce IP lysis buffer. The bound proteins were eluted from the magnetic beads by adding 50 µl of elution buffer (5% SDS, 50 mM TEAB, pH7.55) and heat-denatured for 5 min at 90°C. Light (anti-IgG) and heavy (anti- ZC3H11A) eluted proteins were mixed 1:1.

For the transient overexpression experiments, HeLa cells were transfected with GFP-ZC3H11A(wt) or GFP-ZC3H11A(dZF) expressing plasmids . About 24 h post-transfection cells were lysed in 500 μl of Pierce IP lysis buffer (Thermo Fisher Scientific) supplemented with Halt protease inhibitors and Benzonase nuclease (Sigma, final concentration 25 U/ml). GFP-tagged proteins were immunopurified with GFP-Trap magnetic beads (Chromotek) for 1 h at 4°C. Beads were washed four times in Pierce IP lysis buffer (containing Halt protease inhibitors). Proteins were eluted from the magnetic beads by adding 50 µl of elution buffer (5% SDS, 50mM TEAB, pH 7.55) and heat denaturation for 5 min at 90 °C. Eluted proteins from either the SILAC co-IP or the GFP co-IP were treated by TCEP (5 mM) to reduce disulfide bonds, followed by adding methyl methanethiosulfonate (MMTS) to a final concentration of 15 mM to alkylate cysteines. After that, the lysate was acidified by adding phosphoric acid to a final concentration of 1.2%. The acidified lysate was added to an S-Trap microcolumn (Protifi, Huntington, NY) containing 300 µl of S-Trap buffer (90% MeOH, 100 mM TEAB, pH 7.5) and centrifuged at 4,000 × *g* for 2 min. The S-Trap microcolumn was washed twice with S-Trap buffer. The columns were transferred to new tubes and incubated with 10 ng/μl sequencing-grade trypsin (Promega) (49). The digested proteins were eluted by centrifugation at 4,000 × *g* for 1 min with 50 mM TEAB, 0.2% formic acid (FA), followed by 50% acetonitrile (ACN)/0.2% FA, and finally 80% ACN/0.1% FA. Peptides were dried in a vacuum centrifuge (Thermo Savant SPD SpeedVac, Thermo Fisher Scientific) and dissolved in 1% (v/v) FA. Label-free co-IP of ZC3H11A-Flag was performed by transfecting HeLa cells grown on 100 mm tissue plates with pcDNA3 (control) or with pcDNA-ZC3H11A(wt)- Flag plasmids. About 24 h post-transfection cells were lysed in 500 μl of Pierce IP lysis buffer supplemented with Halt protease inhibitors and Benzonase nuclease (Sigma, final concentration 25 U/ml). Flag-tagged ZC3H11A was immunopurified with anti-Flag(M2) magnetic beads (Sigma) overnight at 4°C. Beads were washed four times in lysis buffer (lacking Benzonase and protease inhibitors). Proteins were eluted from the magnetic beads using 4X Laemmli sample buffer. Eluted proteins were loaded on to a 4-20% Mini-PROTEAN TGX precast gel (BioRad) and run approximately 2 cm in. The gel lanes were cut into two pieces (fractions), and proteins were reduced in-gel with 10 mM DTT in 25 mM NH_4_HCO_3_ (ABC), and thereafter alkylated with 55 mM iodoacetamide in 25 mM ABC, and finally digested with 17 ng/μl sequencing-grade trypsin (Promega) in 25 mM ABC, using a slightly modified in-gel digestion protocol (49). Generated peptides were eluted from the gel pieces using 1% (v/v) FA in 60% (v/v) ACN, dried down in a vacuum centrifuge (Thermo Savant SPD SpeedVac), and finally dissolved in 1% (v/v) FA. All peptides were desalted using Stage Tips (Thermo Fisher Scientific) using the instructions from the manufacturer, and subsequently dissolved in 0.1% (v/v) FA in H_2_O (solvent A) prior to LC-MS analyses.

### Liquid chromatography and mass spectrometric analysis

The desalted peptides were separated on an EASY-nLC 1000 UHPLC (Thermo Fisher Scientific) coupled to an Acclaim PepMap 100 column (2 cm x 75 μm, 3 μm particle size, Thermo Fisher Scientific) coupled in line with an EASY-Spray PepMap RSLC C18 reversed-phase column (50 cm x 75 μm, 2 μm particle size, Thermo Fisher Scientific). The column was heated to 35°C and equilibrated with solvent A. A 2-40% gradient of solvent B (0.1% (v/v) FA in ACN) was run at 250 nl/min for 3 hours. Eluted peptides were injected and analyzed on an Orbitrap Fusion Tribrid mass spectrometer (Thermo Fisher Scientific) controlled by the Xcalibur software (version 4.4.16.14, Thermo Fisher Scientific) and run at Top Speed data-dependent acquisition mode, a spray voltage of 2.4 kV, and an ion-transfer tube temperature of 275°C. MS full scan spectra (m/z 400-2000) at a resolution of 120,000 at m/z 200 were collected in profile mode and analyzed in the Orbitrap with an automatic gain control target (AGC) of 2.0e5, and a maximum injection time of 100 ms. Peptides with an intensity above 5.0e3 were selected for fragmentation utilizing collision-induced dissociation (CID) at a collision energy setting of 30%. Peptide fragments were analyzed in the linear ion trap with an AGC of 1.0e4, and a maximum injection time of 40 ms, with the data acquired in centroid mode. A dynamic exclusion period of the peptides chosen for fragmentation was set to 60 s. Quality control of the instrument setup was monitored using the Promega 6x5 LC-MS/MS Peptide Reference Mix (Promega) and analyzed with the PReMiS software (version 1.0.5.1, Promega).

### Mass spectrometric data analysis

Analysis of mass spectrometric raw files was performed using the MaxQuant quantitative proteomics software package (version 2.1.0.0), including the Andromeda search engine (50, 51). The search was performed with cysteine methyl methanethiosulfonate (MMTS, SILAC co-IP and GFP co-IP) or cysteine carbamidomethylation (label-free co-IP) as a static modification and methionine oxidation and protein N-terminal acetylation as variable modifications. MS1 Orbitrap tolerance was set to 20 ppm, and MS2 ion trap tolerance was set to 0.5 Da. Match between runs enabled identifying peptides where only MS1 data was available in one of the two fractions of each sample for the label-free co-IP, and re-quantify was set to on for the SILAC experiment, in case only one of the partners of a label pair was detected. Peaks were searched against the UniProtKB/Swiss-Prot *Homo sapiens* proteome database (UP000005640, version 2022-04-28), and a maximum of two trypsin mis-cleavages per peptide was allowed. The MaxQuant potential contaminants database was also searched. A decoy search was made against the reversed database, with both peptide and protein false discovery rates set to 1%. Proteins identified with at least two peptides of at least seven amino acids in length were considered reliable.

The MaxQuant output was filtered by removing reversed database hits, proteins only identified by site, and potential contaminants. Proteins with more than one missing value in the three replicates were also filtered out. Normalization of LFQ intensity values was performed using the variance stabilization transformation (VSN) method (52), followed by imputation of missing values using the deterministic minimal value (MinDet) approach (53). The normalized intensities were fitted to a linear model. The significance of differentially enriched (DE) proteins was calculated using empirical Bayes moderated *t*-statistics, utilizing the DEP package for Bioconductor and R (version 1.6.0). *P*-values of the DE proteins were corrected for multiple testing using the Benjamini-Hochberg method. The mass spectrometry proteomics data have been deposited to the ProteomeXchange Consortium (http://proteomecentral.proteomexchange.org) via the PRIDE partner repository (54).

### Immunofluorescence assays

HeLa cells were seeded on coverslips and transfected with 0.25 μg of plasmid DNA. Cells were fixed in 4% paraformaldehyde for 10 min and permeabilized with 0.1% Triton X-100 in PBST (PBS + 0.01% Tween 20) for 10 min at room temperature. The cells on coverslips were blocked with blocking solution (2% BSA/PBST) for 30 min and incubated with the primary antibodies anti-Flag (mouse, Sigma, M2, F1804), anti-PABPN1 (rabbit, Abcam, ab75855), anti- SC35 (mouse, Abcam, ab11826) diluted in the blocking solution overnight at 4°C. Proteins were visualized with the FITC- and TRITC-conjugated secondary antibodies (Sigma, F6005, and T5393) for 1 h at room temperature. Nuclei were stained with DAPI supplemented Fluoromount-G mounting media (Thermo Fisher Scientific). Cells were visualized with a fluorescence microscope (Nikon eclipse 90i), and the images were analyzed with the NIS-elements (Nikon) software.

### RNA crosslinking-immunoprecipitations followed by RT-qPCR (CLIP-qPCR)

The CLIP-qPCR experiments were performed as described (55), with minor modifications. Shortly, transfected A549 cells were washed once with ice-cold 1X PBS and irradiated 2 times with 400 mJ/cm^2^ (254nm) in CL-1000 Ultraviolet Crosslinker (UVP). Quantitative reverse transcription PCR (qRT-PCR) reactions were performed using HOT FIREPOL EvaGreen Supermix (Solis BioDyne) in a QuantStudio 6 Flex Real-Time PCR System (Applied Biosystems). Every sample was analyzed in triplicate. Relative ZC3H11A binding to mRNA is shown after normalizing the immunoprecipitated reactions to the input. Used primer sequences are available in Supplementary Table S1.

### Infectious virus determination

A549 cells were transfected with siScr, siZC3H11A or siPABPN1 for 36 h, followed by HAdV-NanoLuc virus infection (MOI=1). Infected cells were collected at 24 h post-infection (hpi), lyzed in 100 µl of 0.1 M Tris-HCl (pH 8.0), freeze-thawed three times and centrifuged to separate cell supernatant and cell debris. Equal volumes of the cell supernatant were used to re-infect fresh HeLa cells in a 96-well plate for 1 h. Cells were lysed 24 hpi in Passive Lysis Buffer (Promega), and NanoLuc luciferase reporter gene expression was detected using a Nano-Glo® Luciferase Assay System (Promega) and analyzed on a Infinite M200 plate-reader (Tecan).

### RNA isolation and qRT-PCR

Total RNA was isolated using TRIreagent (Sigma) and cleaned with the RapidOut DNA removal kit (Thermo Fischer Scientific). About 1 µg of RNA was used for cDNA synthesis using Maxima H minus reverse transcriptase (Thermo Fischer Scientific) and random primers. qRT-PCR reactions were performed as described above. Every sample was run in triplicate and normalized against the expression of the housekeeping gene HRPT1. Detailed sequences of the used viral and cellular primers are available in Supplementary Table S1.

### TVN-PAT and ePAT experiments

HeLa cells were transfected with indicated siRNAs for 36 h, followed by HAdV-5 infection for an additional 24 h. Total RNA was isolated using TRIreagent (Sigma), DNaseI-treated (Thermo Fischer Scientific), and subjected to TVN-PAT and ePAT analysis essentially as described (38). Gene-specific primers and universal primers for TVN-PAT and ePAT reactions are described in Supplementary Table S1. Semi-quantitative PCR amplifications were done using Phire Taq DNA polymerase (Thermo Fisher Scientific). PCR amplicons were separated on 2% Ultrapure agarose in 1X TAE buffer (Thermo Fischer Scientific) and were visualized with GelRed (Biovitium).

## Data availability

The MS datasets are deposited into a publicly accessible repository.

## Supporting information

This article contains supporting information; Supplementary Figure S1, Supplementary Figure S2, Supplementary Table S1.

## Acknowledgments

We thank Dr. Anette Carlsson for excellent technical support and the rest of the “ZC3H11A” group members for valuable discussions.

## Author contributions

T.P., L.A., and S.Y. conceived the study; K.K., E.S., Z.H., M.L., T.P., and S.Y. performed research; K.K., E.S., Z.H., M.L., T.P., and S.Y. analyzed data; K.K., M.L., G.A., L.A., T.P., and S.Y wrote the paper.

## Funding and additional information

The project was funded by the Knut and Alice Wallenberg Foundation (KAW 2017.0071).

## Competing interests

The authors declare that they have no conflicts of interest with the contents of this article.

---

**Figure S1.**
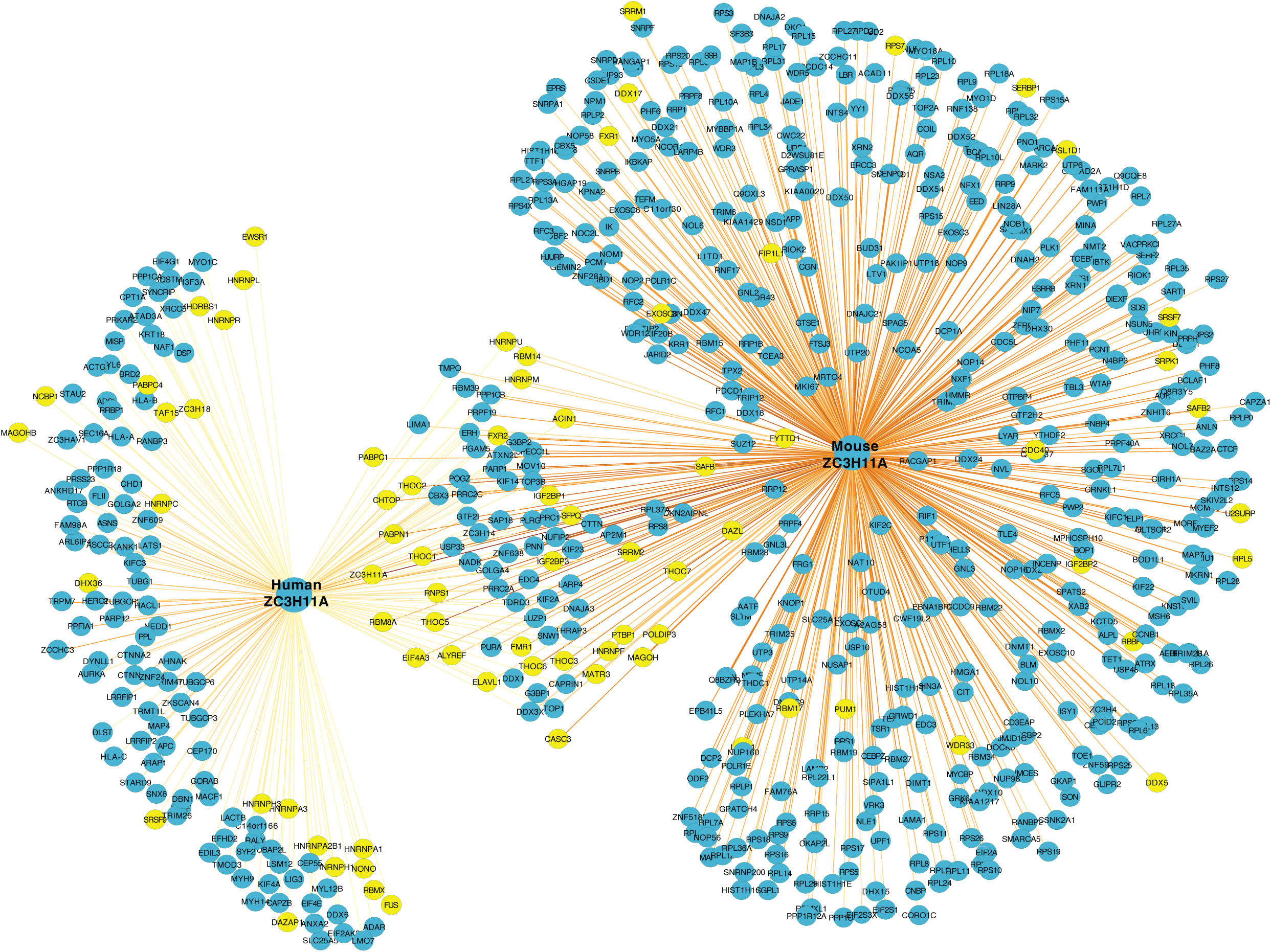
A comparison between human and mouse ZC3H11A interactome. Network analysis of the identified ZC3H11A-interacting partners ZC3H11A in human (HeLa) and mouse (mESCs) cells based on co-Ips (adjusted P <0.05). Yellow circles highlight proteins involved in 3’UTR mRNA processing based on the STRING database.

**Figure S2.**
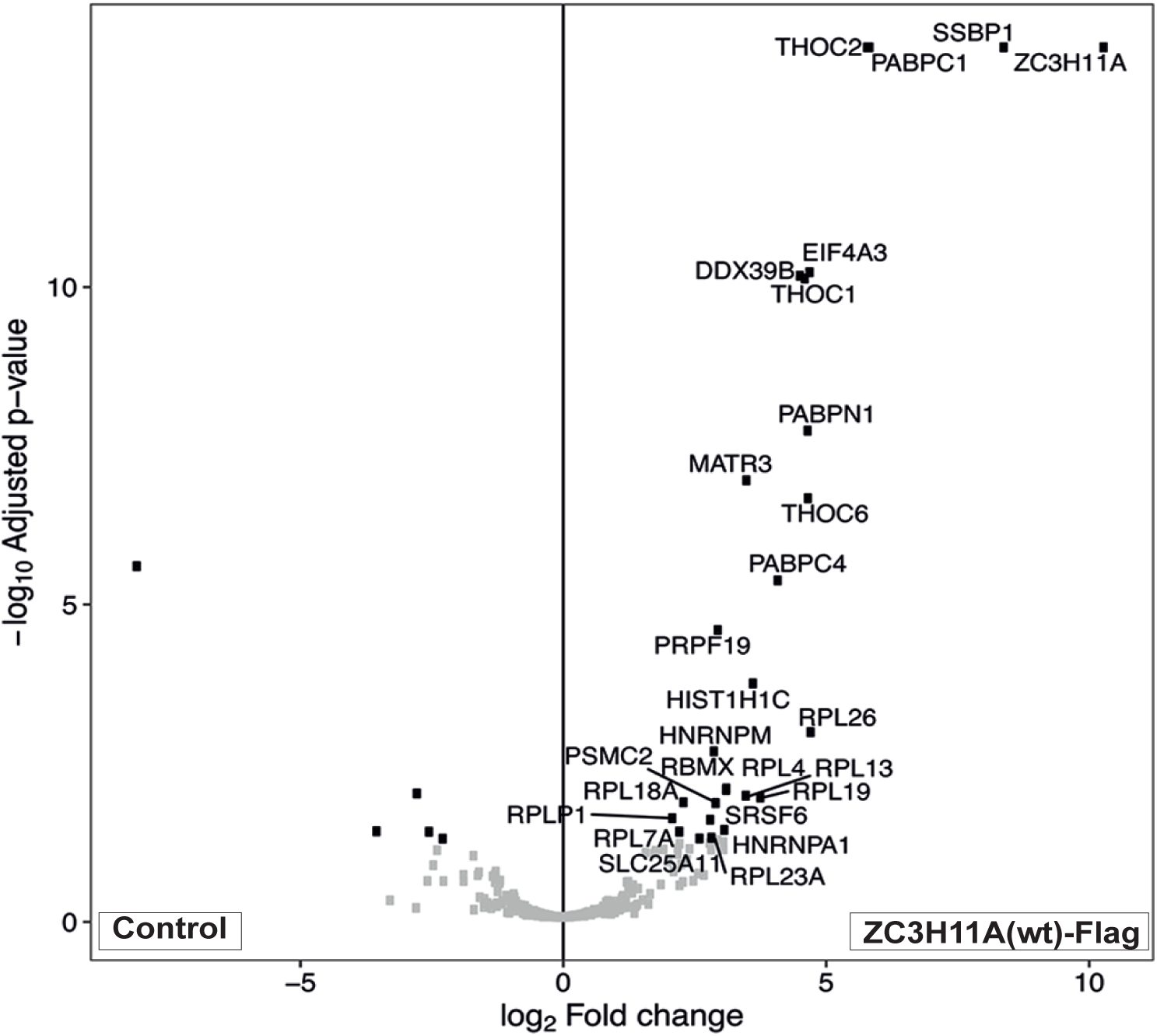
The ZC3H11A-Flag interacts with subunits of the TREX complex and PABPs. Volcano plot showing identified proteins that specifically interact with the ZC3H11A(wt)-Flag protein in HeLa cells. An adjusted P-value cut off of 0.05 and a log2 fold change cut off of 2 was used. Data are shown from a biological triplicate experiment. Control; cells transfected with pcDNA(C-Flag) plasmid, ZC3H11A(wt)-Flag; cells transfected with pcDNA.

**Table S1.**
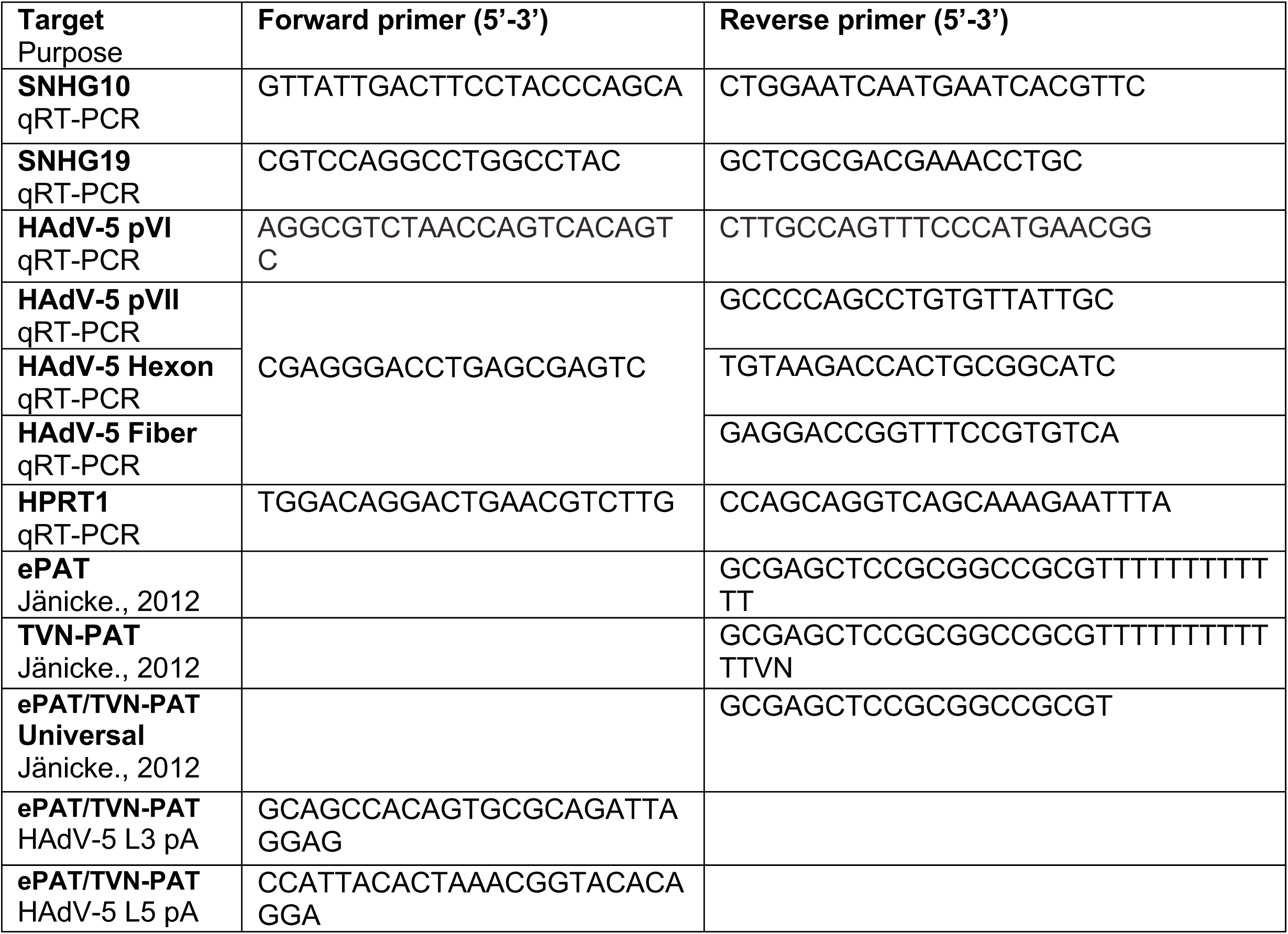
List of PCR primers used in this study

## Notes

### Competing Interest Statement

The authors have declared no competing interest.

